# Population genomic time series data of a natural population suggests adaptive tracking of environmental changes

**DOI:** 10.1101/2020.06.16.154054

**Authors:** Markus Pfenninger, Quentin Foucault

## Abstract

Natural populations are constantly exposed to fluctuating environmental changes that negatively affect their fitness in unpredictable ways. While theoretical models show the possibility of counteracting these environmental changes through rapid evolutionary adaptations, there have been few empirical studies demonstrating such adaptive tracking in natural populations.

Here, we analysed environmental data, fitness-related phenotyping and genomic time-series data sampled over three years from a natural *Chironomus riparius* (Diptera, Insecta) population to address this question. We show that the population’s environment varied significantly on the time scale of the sampling in many selectively relevant dimensions, independently of each other. Similarly, phenotypic fitness components evolved significantly on the same temporal scale (mean 0.32 Haldanes), likewise independent from each other. The allele frequencies of 367,446 SNPs across the genome showed evidence of positive selection. Using temporal correlation of spatially coherent allele frequency changes revealed 35,574 haplotypes with more than one selected SNP. The mean selection coefficient for these haplotypes was 0.30 (s.d. = 0.68). The frequency changes of these haplotypes clustered in 46 different temporal patterns, indicating concerted, independent evolution of many polygenic traits. Nine of these patterns were strongly correlated with measured environmental variables.

Thus, our results suggest that the natural population of *C. riparius* tracks environmental change through rapid polygenic adaptation in many independent dimensions. This is further evidence that natural selection is pervasive at the genomic level and that evolutionary and ecological time scales may not differ at all, at least in some organisms.

## Introduction

Most natural environments consist of an almost infinite number of aspects and parameters that are constantly changing in space and time (Kingsolver *et al*., 2012). Organisms in such habitats therefore potentially have to deal with a fluctuating selective regime (Bell, 2010) where the direction of selection frequently changes (Siepielski *et al*., 2009). The question is whether and to what extent natural populations rapidly and constantly adapt to the fluctuations in their environment in a process termed adaptive tracking (Vander Wal *et al*., 2013). The answer is crucial for our understanding of the relative roles of adaptation vs. phenotypic plasticity (King & Hadfield, 2019), eco-evolutionary processes (Rudman *et al*., 2018), the maintenance of genetic variation (Barton & Keightley, 2002) and the process of balancing selection in natural populations (Abdul-Rahman *et al*., 2021).

Theoretical work has shown that rapid adaptation, a prerequisite for adaptive tracking, is possible almost in real time in natural populations (Messer & Petrov, 2013; Botero *et al*., 2015; Matuszewski *et al*., 2015), in particular if polygenic traits are the target of selection (Jain & Stephan, 2017; Barghi *et al*., 2020). However, despite the importance of the question whether and to what extent selective tracking plays a role in nature, empirical studies are rare. There are a few examples of selective tracking of the fluctuating environment for phenotypic traits (Grant & Grant, 1989; Marrot *et al*., 2017; de Villemereuil *et al*., 2020). In particular the selectively driven beak variability of Galapagos finches in response to different weather conditions in different years, is the classic example of very rapid evolutionary adaptation to a variable environment (Boag & Grant, 1981). On the molecular level, there are several demonstrations of rapid selectively driven changes (Yang *et al*., 2016; Margres *et al*., 2017; Bitter *et al*., 2019; Zong *et al*., 2021). Seasonally selected polymorphisms with correlated allele frequency trajectories were also observed in natural populations of a dipteran species, *Drosophila melanogaster* (Bergland *et al*., 2014; Croze *et al*., 2017). However, studies bringing together the observed temporal fitness differences among different phenotypes with the underlying molecular variants are scarce. In a complex semi-natural experiment (Rudman *et al*., 2021), could demonstrate adaptive tracking on both the phenotypic and genomic level. However, it has been stressed that real-world examples from natural populations are needed (Hendry, 2019).

In a recent article Pfenninger et al. (2022) could show with population genomic time series data and experiments that a natural population of the non-biting midge *Chironomus riparius* (Meigen, 1803) selectively tracked a short-term weather event. Moreover, long term genomic time series data suggested that the spatially unlinked alleles involved in the investigated polygenic trait under selection changed temporally in a correlated fashion over a longer period (Pfenninger *et al*., 2022). This raised the question whether there are more spatially unlinked haplotypes in the genome that change over time in correlated fashion, thus indicating independent responses to the respectively same selective forces. Or, in other words, does this species generally adaptively track the environmental changes in its habitat? To demonstrate adaptive tracking, it is necessary to show that fitness-related phenotypic and selection driven genomic changes happen on the same time scale as selectively relevant environmental changes.

Here, we used environmental data, fitness related phenotyping and population genomic time series data of the same natural *C. riparius* population to tackle this question. *Chironomus riparius* is a multivoltine species with up to 15 generations per year in Europe (Oppold *et al*., 2016). Therefore, the different generations are subjected to widely varying environmental conditions. Accordingly, extensive research on temperature and photoperiod has shown that several traits can and do adapt locally (Waldvogel *et al*., 2018), and temporally among seasons (Foucault *et al*., 2018; Doria *et al*., 2022a). But also other factors are known to act as selection pressures on this species (e.g. organic load, Kraak *et al*. (2000), conductivity, (Pfenninger & Nowak 2008), nitrogen, (Nemec *et al*. 2012), temperature, (Nemec *et al*. 2013) and anthropogenic substances, (Nowak *et al*. 2009). The high effective and demographic population size (> 1,000,000, Waldvogel *et al*. (2018)) and the very high number of offspring per breeding pair (400-800) allows for rapid adaptation (Pfenninger & Foucault, 2020). A range of genomic resources and parameters are available (Schmidt *et al*. 2020; Oppold & Pfenninger 2017) and established routines for evolutionary experiments in the laboratory exist (Foucault *et al*., 2019b), which makes the species a suitable object to test the hypothesis that adaptive tracking takes place in natural populations. Here, we addressed the following questions:

- Did known selection pressures change over the periods monitored in the natural population, thus providing a fluctuating selective regime?
- Is there fitness-related phenotypic evolution on the same time scale in the population?
- Can we find evidence for selection-driven allele frequency changes in the genome?
- Are there spatially unlinked selected haplotypes whose allele frequencies evolve in a temporally correlated fashion, indicative of polygenic adaptation?
- If so, how many of such selected trajectory clusters can we identify?
- Are the frequency changes of the haplotypes in these clusters correlated with recorded environmental changes?

## Material and Methods

### Sampling

In February 2016, more than 500 living red *thummi*-type chironomid larvae were sampled with a sieve at a single site situated in a small river (Hasselbach, Hessen, Germany 50.167562°N, 9.083542°E) following the protocol of (Foucault *et al*., 2019b). The specimen’s specific identity was ascertained by DNA-barcoding of a mitochondrial (COI) and a nuclear locus (L44). Eighty thus identified *C. riparius* were pooled and subjected to PoolSeq sequencing (see below). The sampling was likewise repeated in September 2016, assuming that the two annual samples thus were subjected to the selective extremes of the winter and summer season, respectively. This sampling scheme was repeated for the two following years, 2017 and 2018, resulting in a time series of six population pools (NP0-NP5). From February to September, the population produced about seven generations, in the period from autumn to early spring about three. The entire sampling period thus covered about 27 generations (Suppl. Fig. 1). We used additional *C. riparius* individuals from the field, sampled at the same time as the individuals for genomic analysis except for September 2018, to perform standardised life cycle fitness tests at three temperatures (14°C, 20°C and 26°C).

### Environmental parameters

Mean monthly water parameters for the period 2009-2018 were obtained from the Hasselbach water treatment plant of the Abwasserverband Freigericht. The values were continuously measured automatically in the river past the outflow, less than 50 m from the sampling site. The following parameters were recorded: water temperature [°C], conductivity [mS/cm], oxygen, biological oxygen demand (BOD), chemical oxygen demand (COD), NH_4_, NO_3_, NO_2_, N_tot_ and PO_4,_ all [mg/l]. To assess the similarity of the entirety of the parameters over time, the monthly means were summarised in a seasonal mean for the period before sampling, *i*.*e*. from March to September and from October to February. These values were subjected to a Principal Components Analysis (PCA). Temporal autocorrelation was calculated in Past 4 (Hammer *et al*., 2001).

### Phenotypic evolution

We used an established life cycle fitness test to assess three fitness component phenotypes of the natural population under standard conditions in a common garden experiment. The experiment was carried out at 14°C which corresponds to the annual average water temperature at the sampling site. A fraction of the sampled larvae from each sampling date was divided into five replicates (30 individuals each) and brought to hatch and reproduce. To distinguish phenotypic evolution from phenotypic plasticity, the resulting F1 generation was monitored for the parameters developmental rate (measured as emergence mean time 50 EmT50), fecundity (viable eggs per female), mortality (proportion of emerging midges) and a resulting overall fitness parameter (population growth rate per day). The assessed phenotypes are known to have a complex genetic basis (Doria *et al*., 2022a). We tested with an ANOVA for significant changes in mean trait values. A detailed description of the life cycle fitness test can be found in (Foucault *et al*., 2019b).

We calculated rates of phenotypic evolution among sampling dates for each trait in Haldanes (h) as:

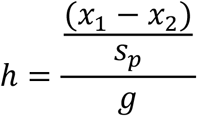

where x_n_ represent the mean trait value at the time point n, s_p_ the pooled standard deviation and g the number of generations between the two time points (Haldane, 1949)

### Population genomic analyses of pooled samples

DNA was extracted for each pool from the field or the experiment using the Qiagen blood and tissue extraction kit on pooled samples of 80 larval head capsules, respectively. Integrity and quality of extracted DNA was controlled using electrophoresis, and the DNA concentration for each samples measured with a Qubit fluorimeter (Invitrogen).

Whole genome pool-sequencing was carried out on an Illumina MySeq with 250bp paired end reads. Reads were trimmed using the wrapper tool autotrim (Waldvogel *et al*., 2018) that integrates trimmomatic (Bolger *et al*., 2014) for trimming and fastQC (Andrews 2010) for quality control. The trimmed reads were then mapped on the latest *C. riparius* reference genome (Schmidt *et al*., 2020) using the BWA mem algorithm (Li & Durbin, 2009). Low quality reads were subsequently filtered and SNPs were initially called using samtools(Li *et al*., 2009). The pipelines Popoolation and Popoolation2 (Kofler *et al*., 2011a, 2011b) were used to call SNPs, remove indels, and to estimate genetic diversity as Watterson’s theta each pool. Allele frequencies for all SNPs with coverage between 15x and 70x were estimated with the R library PoolSeq (Taus *et al*., 2017).

### Identification of allele frequency trajectories with signs of selection

Selected SNP loci were identified by allele-frequency changes (AFC) larger than expected by neutral drift. First, Fisher’s exact tests for the AFCs among all consecutive time points (NP0 > NP1 etc.) and among the first and last sampling were calculated. This test identifies AFC larger than expected by chance, but does not account for genetic drift expected. Therefore, neutral simulations were used to compute false discovery rate q-values < 0.001 with parameters (number of SNPs, starting allele frequencies in the ancestral population, sequence coverage, number of generations) matching those of the respective sample. As effective population size, we used 15,000, which constitutes a very conservative estimate (see (Waldvogel *et al*., 2018)). All calculations and simulations were performed with the R-library poolSeq (Taus *et al*., 2017). A SNP locus was considered showing signs of selection if at least one sampling interval or between the first and the last sample period had an AFC significantly larger than expected. The allele frequency trajectory, i.e. the vector of allele frequencies of such putatively selected SNP site over time was recorded.

### Inference of genome wide level of Linkage Disequilibrium

The expected length of segregating haplotypes depends on the recombination rate and their age. The former can be approximated by an estimate of linkage disequilibrium (LD,(Feder *et al*., 2012). We estimated the extent of LD and its decay using the maximum likelihood method of (Feder *et al*., 2012) on one of the pools (NP0). This method exploits the haplotype information contained in (paired) reads and has been shown to yield accurate estimates if mean LD decay to background levels is shorter than the read length (Feder *et al*., 2012).

### Clustering of spatially coherent SNPs into haplotypes

Most SNPs showing larger than expected AFC are hitchhiking with the actually selected site in selected haplotypes. The extent of such haplotypes was determined following the rationale of (Franssen *et al*., 2016) that physically linked variants following the same evolutionary trajectory should show correlated allele frequencies through time. However, contrary to an experimental evolution situation with a constant selection pressure, it could not be expected that the observed AFC were always in the same direction. Since this is a prerequisite for the software used in Franssen *et al*. (2016), we had to develop our own algorithm. Our algorithm considered consecutively all temporal trajectories of significant SNPs along a scaffold. A SNP was added to a haplotype if the absolute correlation coefficient of its temporal trajectory to any other trajectory of the SNPs already in the haplotype equalled or exceeded a given threshold (0.75). This was repeated until the next SNP position did not meet the criterion. The number of SNPs in the haplotype, its start and end position and the mean allele frequency were then recorded and a new haplotype was started at the next position considered. The analysis was carried out with a custom Python script.

### Validation of haplotypes

Inferred haplotypes were validated with 30 whole genome resequenced individuals (25X coverage, 150 bp paired end reads) from all sampling periods. The reads from these individuals were quality checked, mapped against the reference genome as described above and the genotypes called with the GATK pipeline (Van der Auwera *et al*., 2013). Consensus sequences of one hundred randomly chosen haplotypes with more than 2 SNPs, including one neighbouring haplotype were extracted from the individual VCF-files, aligned and phased with FastPhase in DNAsp (Rozas *et al*., 2017). The resulting phased alignments were checked for linkage of the previously identified target haplotype’s typical alleles and linkage equilibrium to the neighbouring haplotypes in the same software.

### Calculation of selection coefficients for haplotypes

The selection coefficient *s* was calculated from the change in mean allele frequencies between samplings for all selected haplotypes by the formula given in (Kimura, 1970). Assuming codominance (i.e. h = 0.5), the selection coefficient *s* can be calculated as

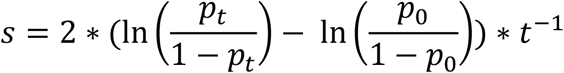

where p_0_ is the allele frequency of the rising allele in the ancestral population and p_t_ its frequency in generation t.

### Hierarchical clustering of mean haplotype trajectories

For each haplotype, we calculated the mean allele frequency trajectory (MHT) over the allele frequencies of all constituting SNPs. The matrix of absolute pairwise correlation coefficients among all MHT was transformed into a distance matrix (1 – correlation coefficient), hierarchically clustered with UPGMA and the resulting dendrogram plotted. The number of clusters considered was determined by applying a clustering threshold of 0.25, corresponding to a minimum similarity of 0.75 of all MHT within a cluster.

To determine this threshold sensibly balancing between arbitrary lumping of actually different trajectories and over-splitting of trajectories differing rather by stochastic fluctuations, we applied a randomisation approach. From the empirical AFC distribution of all SNPs in the analysed set (i.e. at least one significant AFC among consecutive sampling periods or between the first and last sampling period, see above), we randomly drew, 1,000 times the empirically found number of MHTs. Starting with a distance threshold of 0.05, we iteratively increased the threshold by 0.05. In each iteration, the trajectories of the 1000 random MHTs were hierarchically clustered as described above and the number of resulting clusters recorded. The simulated distribution was then compared to the empirically observed number of clusters, applying the same threshold. The smallest distance threshold for which the observed number of clusters was smaller than the 0.05 quantile of the randomly simulated distribution was deemed appropriate for the analysis. This was the case for a distance threshold of 0.25; i.e. the trajectories within a cluster showed a correlation of at least 0.75. The analysis was performed with a custom Python script.

### Correlation of environmental parameters with haplotype trajectories

The seasonal mean environmental parameters (see above) before the first and the respective period between two sampling times were correlated to the means of all identified MHT patterns. To avoid problems with multiple testing and lacking statistical power (Sjölander & Vansteelandt, 2019), we employed a Bayesian implementation of a test on correlation with default values (Bååth, 2014). We report only correlations with convincing posterior evidence (>95% posterior credibility) for positive, respectively negative associations.

### GO-term enrichment analysis

The annotated gene overlapping with a haplotype was identified as putative selection target. The resulting gene lists for all member haplotypes of each MHT pattern were tested for over-representation of GO terms in the category “biological function” in the R-library TopGO with a Fisher’s exact test (Alexa & Rahnenführer, 2016).

## Results

### Fluctuating enviroment

Measured physico-chemical water parameters varied substantially through time (Figure 1, Suppl. Fig.2). Comparison with a broken stick model suggested that the first two Principal Component axes were meaningful. Axis 1 (51.8% of variation) of the PCA was mainly determined by parameters describing organic water pollution (chemical oxygen demand, NH_4_, biological oxygen demand, conductivity), axis 2 (21.7%) reflected mostly seasonal changes (temperature, oxygen, PO_4_, Figure 1). The measured environmental regime never returned close to any previous state. This was mirrored in the poor temporal autocorrelation of the parameters, except for the regular seasonal temperature changes (Suppl. Fig. 3).

**Figure 1.**
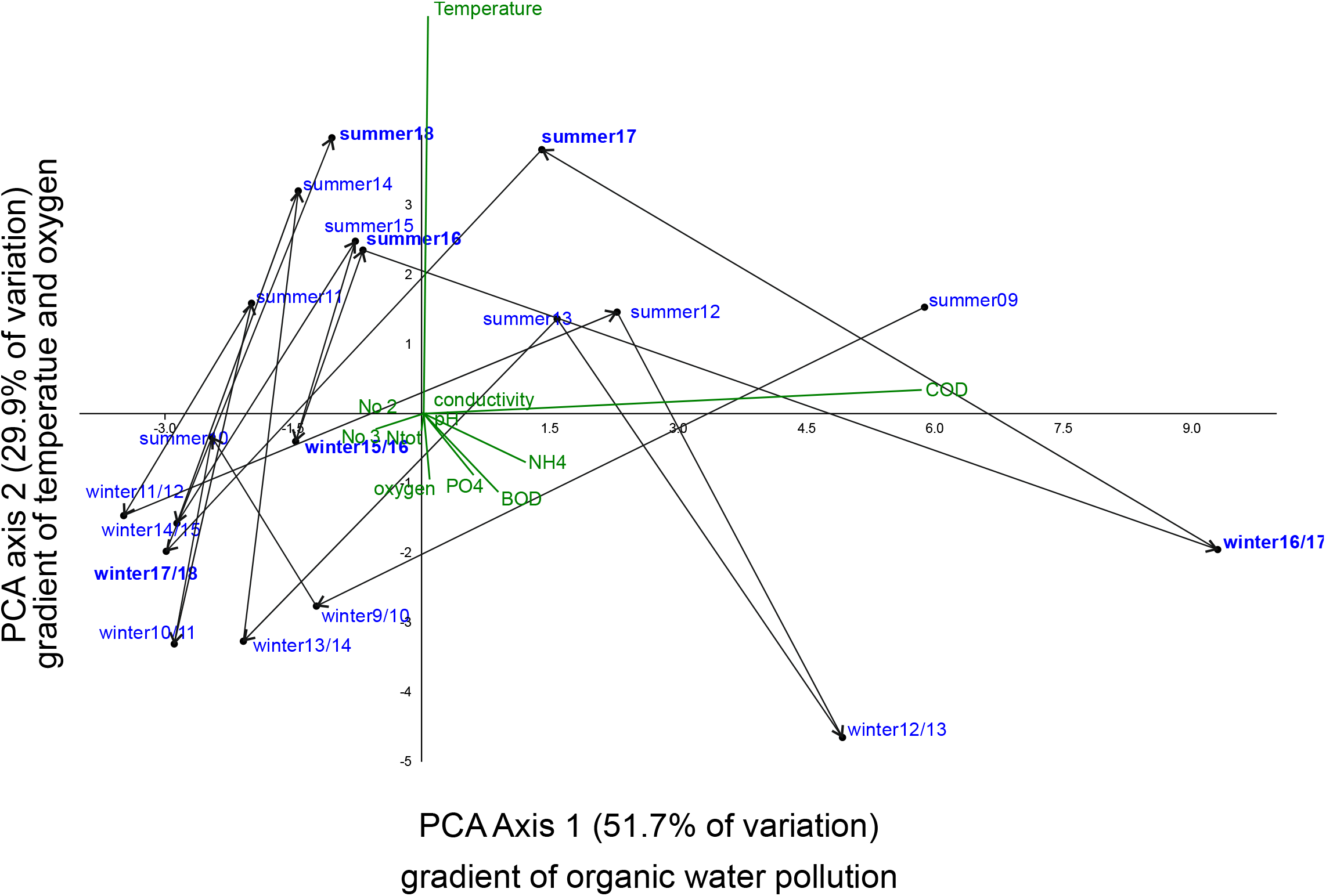
PCA on seasonal physico-chemical water parameters in the natural population for the decade from 2009-2018. Shown are the first two axes, accounting for 82.2% of environmental variation. Periods after which phenotypic and genomic assessments of the population were performed are indicated in bold. The temporal trajectory of the environmental conditions are indicated by arrows connecting consecutive periods.

### Strong phenotypic evolution

Exposed to the same standard conditions in a common garden setting, the F1 generation derived from the natural population at different points in time showed signs of strong and rapid phenotypic evolution, with idiosyncratic patterns across traits. The EmT50 value varied strongly and significantly with the seasons (F = 11,53, d.f. = 4, p = 5 × 10^−5^) between 35.9 and 42.5 days, with an overlaid overall decreasing trend (Figure 2A). Mean evolutionary rate for the trait between sampling dates was 0.29 Haldanes (min = 0.07, max = 0.42) with a reversal of direction between each sampling date (Figure 2A). Mortality varied also significantly, albeit marginally among sampling dates (F = 3.25, d.f. = 4, p = 0.03). Between 11% and 45% of the midges did not hatch in the laboratory environment (Figure 2B). The rate of evolution was 0.22 (min = 0.07, max = 0.31) with two changes of direction (Figure 1B). The strongest phenotypic evolution was observed for fecundity (Figure 2C). The mean fecundity varied significantly between 84 and 286 offspring per female (F = 6.07, d.f. = 4, p = 0.002). The evolutionary rate of change for the trait was on average 0.45 (min = 0.20, max = 0.96) standard deviations per generation. The direction of the evolution changed twice during the period of observation (Figure 2C). The resulting population fitness remained always positive (range = 1.112 – 1.145), but varied significantly among sampling dates (F = 3.84, d.f. = 4, p = 0.02, Figure 1D).

**Figure 2.**
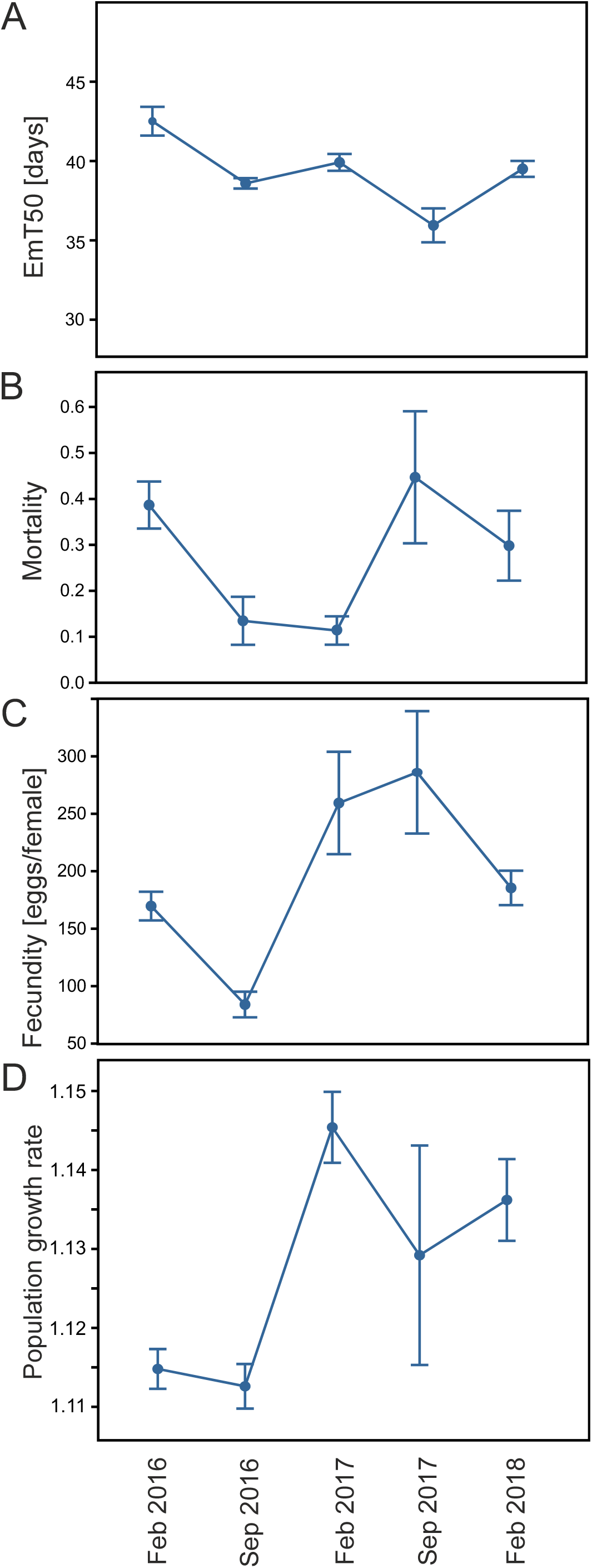
Development of phenotypic traits and population fitness over time in the natural population. A) Developmental time measured as EmT50. B) Mortality measured as proportion of emerged individuals. C) Fecundity as viable eggs per female. D) Population growth rate per day as integrative measure of population fitness.

### Large parts of the genome showed evidence of selective evolution

The genome-wide mean estimate of distance-dependent linkage disequilibrium (LD) dropped to a mean r^2^ of 0.5 within 10 base pairs and to the genome-wide background level of about 0.2 within 120 bp (Suppl. Fig. 5). Consequently, the expected haplotype structure was extremely short.

Allele frequency estimates for all sampling dates were obtained for 22,693,348 SNP positions in the genome. Neutral drift simulations, taking observed sampling variance of the poolSeq approach into account, showed that the chosen selection detection threshold of 2.0 -log_10_p for an allele frequency change (AFC) between consecutive sampling times or between the first and the last sampling implied a false discovery rate of q < 0.001. This corresponded to an AFC of at least 0.15. Under the condition of at least one selected AFC between any two consecutive sampling times or the first and the last sampling time, identified a set of 367,446 SNPs with signs of selection.

Clustering spatially coherent SNPs with signs of selection into haplotypes resulted in 281,153 haplotypes of which 35,574 consisted of more than a single SNP. Of the 100 haplotypes randomly chosen for validation, 98 showed the expected linkage between the marker SNPs of the haplotype (Suppl. Fig. 6). In two cases, both with more than 2 haplotype marker SNPs, one of the marginal SNPs was only slightly linked (r < 0.6). All haplotypes checked were distinct from their respective spatial neighbours. Only the 35,574 haplotypes with more than a single SNP were retained for subsequent analyses. In these, the average number of SNPs per haplotype was 3.30 +/- 2.64 (mean +/- s.d.). The mean length of inferred haplotypes was 893.1 +/- 1790.3 bp. Neighbouring haplotypes on the same scaffold were on average 3466.0 +/- 4293.5 bp apart. Of all 13,490 annotated genes in the genome, 5793 (= 42.9%) harboured at least one selected haplotype, some more.

The mean selection coefficient over all retained haplotypes was 0.30 with a standard deviation of 0.68. The latter is equivalent to the intensity of selection over time, and its much larger value relative to the mean indicated large changes in magnitude and frequent reversals in the direction of selection (Bell 2010).

### Clustering of haplotype trajectories

To avoid using an arbitrary clustering threshold, we used a Monte-Carlo simulation approach. Clustering of simulated random MHTs, drawn from the observed AFC distribution, indicated that a clustering threshold of 0.75 yielded informative clusters without undue lumping (Suppl. Fig. 7). Applying this threshold, the 35,574 MHTs were grouped into 46 distinct cluster (Fig. 2). The number of MHT cluster members was highly different. The largest cluster (MHT16) contained 17,914 haplotype trajectories, the smallest merely three (MHT28), the mean was 816.8 (Suppl. Table 2).

### Correlation of haplotype trajectories to environmental parameter changes and GO analysis

Nine MHT cluster patterns were highly correlated to measured water parameters in the period before sampling. Three MHTs were correlated to changes in NO_3_. One MHTs followed variation on Biological Oxygen Demand (BOD) closely. Two other MHTs were associated to NO_2_, MHT3 was correlated to PO_4_, MHT 38 to temperature, while the strongest association was found for MHT 34 and oxygen (Figure 3). For 44 of the 46 MHT clusters, at least one significantly enriched GO term was found (Suppl. Table 2). Most enriched terms for the individual MHT clusters were on a very high hierarchical level. Consequently, they were rather uninformative or affected only a small minority of the genes with overlap to a selected haplotype. However, some hinted at a specific biological process or supported the observed association of the MHT cluster to a measured physico-chemical variable.

**Figure 3.**
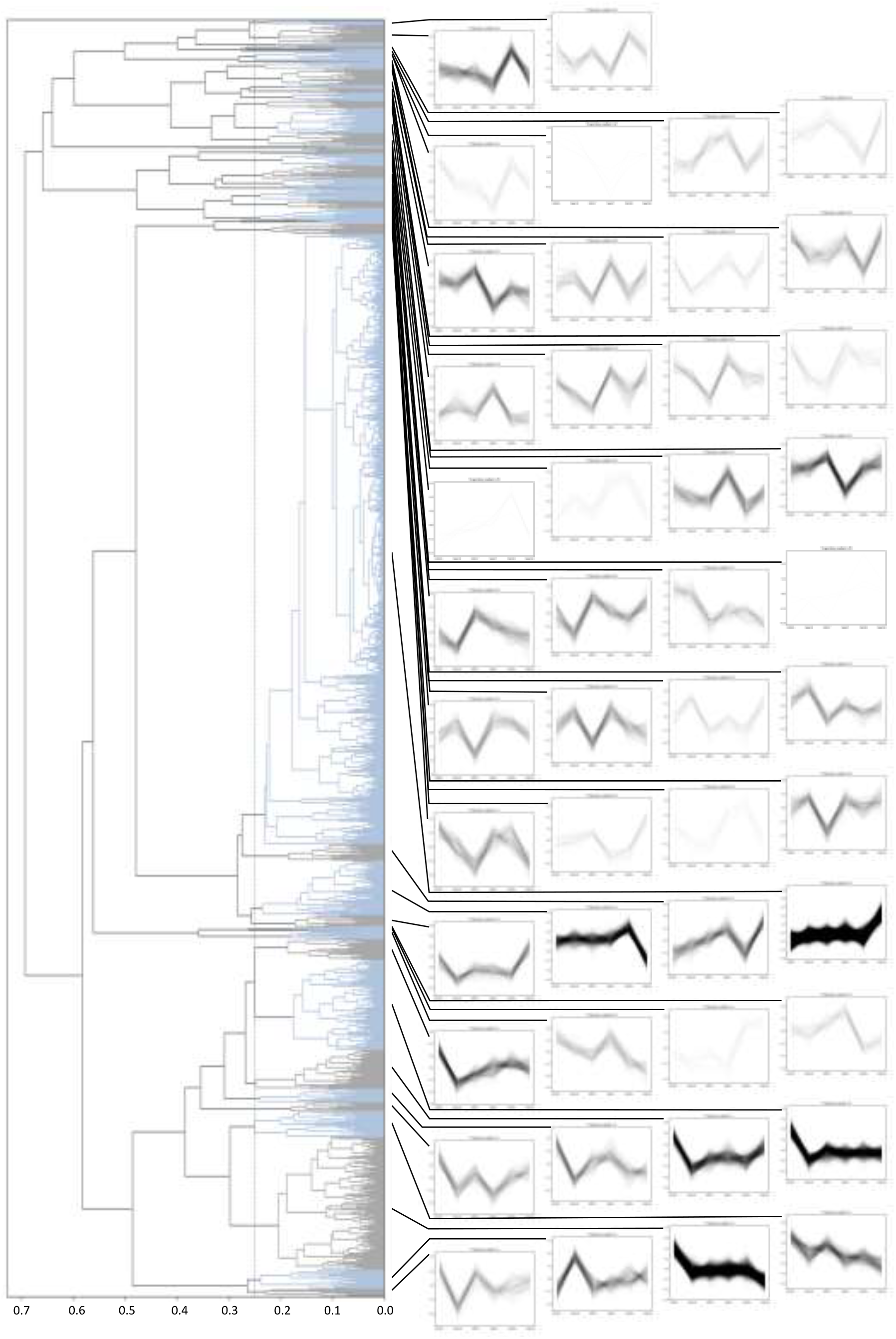
Temporal dynamics and complexity of allele frequencies trajectories with signs of selection. The left side of the panel shows a UPGMA-dendrogram on the pairwise distance (1 - correlation coefficients) of all 35,574 haplotype MHTs with at least one significant AFC. The hatched vertical line presents the inferred clustering threshold of 0.25. The right side of the panel shows the MHTs for each cluster. For better visibility and comparability, the individual MHTs in each cluster were polarised into the same direction and normalised by substracting the respective mean.

**Fig 3.**
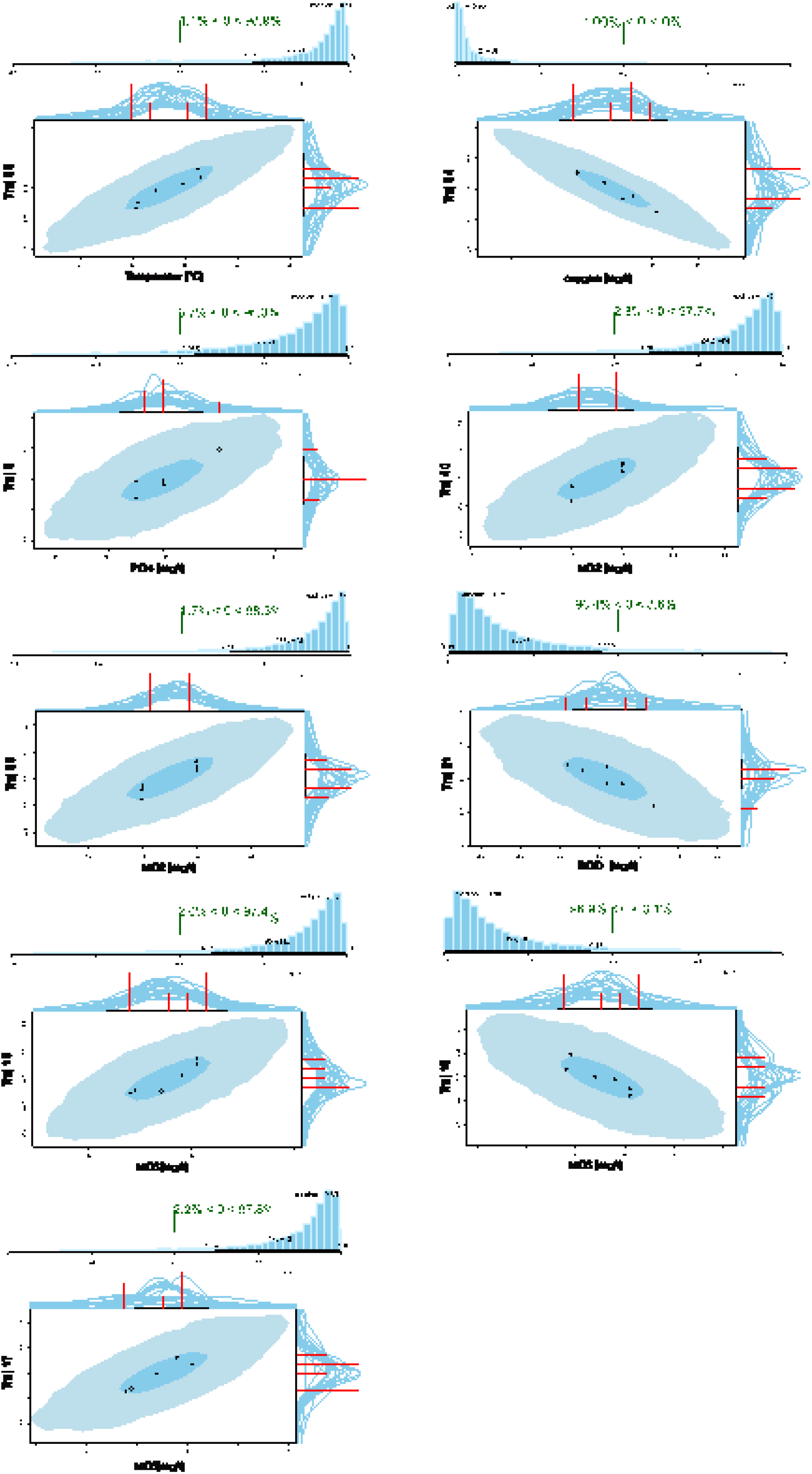
Correlations between measured environmental parameters and MHTs. Shown are only associations with a posterior credibility > 95%. Please note that the direction of the correlation depends on the arbitrary polarisation of the alleles in the allele frequency trajectory.

The enriched terms for MHT4 (GO terms *0051704 multiorganism process, 0006952 defense response*) suggested reaction to a biological interaction. The largest cluster MHT16 contained hundreds of genes involved in osmoregulation, reflected in the enrichment for GO terms *0055085 transmembrane transport* and *0006811 ion transport*. Despite affecting only about 20 genes, MHT36 showed an enrichment of terms associated with *porphyrin-containing compound biosynthetic process (0006779)* and *pigment biosynthetic process (046148)*. Genes affected by MHT40 haplotypes were enriched for signal transduction processes (*0007186 G protein-coupled receptor signaling pathway*), perhaps in conjunction with hormones (*1901362 organic cyclic compound biosynthetic process, 0008202 steroid metabolic process*). MHT3 was enriched for term (*0006811 ion transport*). MHT39, with *calcium (transmembrane) transport* (terms *0006813* and *0071805*), which is co-annotated to *nitric-oxide synthase inhibitor activity*. There was an enrichment of the term *0005975 carbohydrate metabolic process* in MHT21. MHT22 was enriched *for 0007688 sensory perception of smell* and *0006979 response to oxidative stress*.

## Discussion

### Complex and unpredictable multidimensional fluctuating selective regime

The largely uncorrelated temporal changes as well as the observed range of the measured physico-chemical water parameters suggested that the local environment presented a highly fluctuating, multidimensional selective regime for *C. riparius*. For most of these parameters, the fitness relevance and thus their selective potential for the species has already been shown (Ristola *et al*., 1999; Kraak *et al*., 2000; Pfenninger & Nowak, 2008; Nemec *et al*., 2012, 2013). Even though some parameters showed seasonal changes, these were either overlain by long-term trends or were minor in comparison to the observed changes that did not seem to follow any regular pattern. In consequence, the selective regime as a whole defined by this physico-chemical parameter set was at no point in time identical or even similar to any other period surveyed. Moreover, only a small subset of the potentially selectively relevant biotic and abiotic parameters was measured. For example, predation (Correia *et al*., 2005) and other intra- and interspecific ecological interactions that are known to play an important role for the fitness of *C. riparius* (van de Bund & Davids, 1993; Hooper *et al*., 2003) were not taken into account. It is therefore reasonable to assume that an unknown, but much larger number of selective factors is simultaneously, yet in strength and direction more or less independently, acting on this population. The local habitat is therefore probably conforming to the multi-scale theory of the environment (Bell, 2010).

### Continuous phenotypic evolution with frequent changes in direction

The measured phenotypes of the natural population varied significantly among sampling dates and inversed the direction of change repeatedly and independently across traits. To distinguish between phenotypic plasticity and adaptation, we measured the traits in the F1 generation under standard laboratory conditions, which may be viewed as too few to exclude imprinting effects (Glastad *et al*., 2019). The laboratory conditions, however, are themselves not selectively neutral for field populations (Pfenninger & Foucault, 2020). Given the propensity for rapid adaptation in the laboratory, particularly in the first few generations when confronted with a new stressor (e.g. (Doria *et al*., 2022a, 2022b), it was necessary to balance between these two pitfalls.

We observed strong evolutionary responses, with per generation rates that were in range of those observed for insects in laboratory experiments (Hoffmann & Ross, 2018) or in the semi-natural setting of (Rudman *et al*., 2021). Because there were frequent changes in the direction of change, the chosen test interval was crucial; as in the latter study, the choice of another time scale could have suggested no phenotypic change at all.

Most importantly for the diagnosis of adaptive tracking was that phenotypic evolution occurred on the same temporal scale as the environmental change. However, we did not test for a direct link between any environmental parameter and a phenotypic trait, because i) the measured traits are rather highly inclusive fitness components, thus integrating the influence of many lower level traits, ii) transient, short term extreme events were not recorded by the long-term averages, but are known to exert important selective pressure (Pfenninger *et al*., 2022). All measured traits have strong links to fitness, which also varied over the monitored period. This concurred to selective dynamic patterns generally known from natural populations (Kingsolver *et al*., 2012).

### Rapid and pervasive selective genomic evolution

The analysis of population genomic time series showed that tens of thousands loci showed signs of at least one selection-driven change in allele frequency across sampling periods, corresponding to less than 30 generations. Rapid and substantial changes in allele frequencies were observed throughout the genome. As the chosen approach accounted for drift and sampling variance (Taus *et al*., 2017), and immigration was unlikely to change allele frequencies noticeably (Foucault *et al*., 2019a), the observed significant changes are likely due to selection alone. This supports the increasing observations of rapid adaptive change in natural populations (e.g. (Torda *et al*., 2017; Dayan *et al*., 2019). The observation that the standard deviation of the selection coefficients, an equivalent to selection intensity, was much larger than the respective mean (0.68 vs. 0.30) proved substantial changes in magnitude and frequent reversals in the direction of selection. The acting selection regime thus corresponded to the multi-scale model of fluctuating selection advocated by (Bell, 2010).

The propensity to rapid adaptation with simultaneous preservation of genetic variation in *C. riparius* is thus probably due to an interplay of genomic, life cycle and demographic traits of the species. On the genomic level, the low observed mean LD resulted in a highly fragmented haplotype structure, comparable in scale to other insects like *Drosophila* (Feder *et al*., 2012). A short range haplotype structure is a precondition for the observed short distance between haplotypes responding to selection. Since only haplotypes with more than a single SNP were considered, this may even present an underestimate, because ancient selected polymorphisms may very well be completely separated from their background by recombination (Pfenninger *et al*., 2022). In some instances, even different parts of the same gene followed different evolutionary trajectories. The observed fine-scaled selective response to independently fluctuating environmental parameters requires that many different recombinational variants are permanently present in the population. This may be one reason for the gross overproduction of offspring in the species (Foucault *et al*., 2019b). A large offspring number implies a respectively large number of independent recombinational events, creating a wide variety of new genotypes from which to select. Together with the large, fluctuating demographic and effective population size, the preconditions for rapid, fine-scaled and variable evolutionary reaction to fluctuating changes in the selective regime are therefore given.

### The number of temporal evolutionary trajectories is limited but dynamic

In comparison to the huge number of potentially independent loci in terms of physical linkage reacting to selection, the number of statistically distinguishable evolutionary trajectories was quite limited (46). Despite expected trade-offs and pleiotropy between loci (Li *et al*., 2019), this suggested that rather the number of detectable selection pressures was limited than the capacity of the population to follow them. That the majority of genes was not measurably affected by selection suggested unused adaptive potential, but could also be due to small effect size loci that did not reach statistical significance in the present analysis.

While the order of magnitude of estimated effectively acting selection pressures is certainly correct, the exact value may not be. On the one hand, the statistically derived clustering threshold was possibly too rigid and therefore distinguished too many clusters, as indicated by the correlation of some very similar MHT clusters (MHT21, MHT22 and MHT17, MHT18, MHT19) to the same environmental variables (Fig. 3). On the other hand, in longer time series other selection pressures could have elicited additional statistically detectable MHT or split co-varying but distinctive selection pressures into distinguishable haplotype trajectories, thus increasing the estimate of acting selective agents. A hint in this direction was the increase from 36 distinguishable MHTs in a preliminary analysis after five temporal samplings (data not shown) to 46 after six. It was therefore possible that the number of selection pressures the population can effectively follow independently within few generations may be finally limited, but could be different than estimated here. However, general conclusions will require the application of the framework presented here to more populations over more generations and different species.

Changes in intensity and direction of MHTs were observed frequently, thus corroborating the results gained from phenotypic traits. The number of haplotypes that showed a continuous increase or decline in allele frequency over the monitored period were relatively limited (MHT4, 724 loci), suggesting that adaptation proceeds mainly by short time scale tracking of varying selection pressures and not by slow fixation of alleles.

As expected from studies on *Drosophila* (Bergland *et al*., 2014), MHT patterns reflecting seasonal variation were identified. MHT38, highly correlated to temperature and MHT40 showed respective patterns, accounting for about 1.2% of all selected haplotypes. This is in the range found for seasonal fluctuations of allele frequencies in *Drosophila* (Bergland *et al*., 2014; Machado *et al*., 2021). However, compared to the high variability of the remaining MHT patterns, the seasonal variations in temperature, precipitation, resource availability etc. were surprisingly of relatively minor importance.

### Environmental associations and functional gene enrichments of MHTs

The number of MHTs in each cluster was highly different, ranging from a few to many thousands. On average, many independent loci were therefore involved in the same adaptive response, which is typical for polygenic traits (Jain & Stephan, 2017). However, enrichment analyses in the GO category Biological Process for genes overlapping with haplotypes from respective MHT clusters rarely yielded clearly interpretable patterns. In some instances, hypotheses on the acting selection pressures could be gained from the enrichment analyses. In particular MHT4 suggested a biological interaction, perhaps in defence to parasites. Immune genes adaptively tracking the abundance of potential pathogens over seasons were also found in wild *D. melanogaster* populations (Behrman *et al*., 2018). Another interesting set of enriched genes associated with porphyrin containing pigment metabolism was found in MHT36. The most important pigments in Chironomids are hemoglobins, giving the characteristic red colour to these midges (also called “bloodworms”). They play an important role in hypoxia/anoxia (Grazioli *et al*., 2016) and are negatively influenced in their function by nitrite (Ha & Choi, 2008). It was thus not surprising that the highest correlations of this MHT were to oxygen (r = 0.74) and NO_2_ (r = 0.78), respectively. Less than a quarter of the MHT patterns were highly (> 0.9) correlated with the few measured environmental parameters, indicating that only a small proportion of selectively relevant parameters were measured. Some correlations suggestively matched the inferred functions from the GO enrichment analysis. MHT39, enriched for genes involved in *nitric-oxide synthase inhibitor activity*, was highly correlated to NO_2_, underlining the importance of nitrogen metabolism for the species (Pfenninger & Nowak, 2008). Another match was found for MHT21, strongly associated with biological oxygen demand, which reflects the availability of organic matter in the stream, an important food source for the midges (Ristola *et al*., 1999). Among the GO terms enriched in this MHT, *carbohydrate metabolism* was found, insinuating an adaptive reaction to the food availability.

### Adaptive tracking of environmental changes

The observed substantial rapid phenotypic and selectively driven genomic evolution with changing signs on the same time scale as putatively selectively relevant environmental variation strongly suggested that the natural *C. riparius* population studied here adaptively tracked their environment. This implies the existence of abundant, unlinked genetic variation under fluctuating selection. In classical population genetic predictions, adaptive tracking as observed here is viewed as unlikely to evolve, because fluctuating selection should rapidly eliminate genetic variation (Hedrick, 2006). Several recent models, however, have called these predictions into question. It was shown that in a multivoltine species, adaptive tracking is more likely to evolve than phenotypic plasticity or bet-hedging (Botero *et al*., 2015). Dominance shifts of adaptive alleles (Wittmann *et al*., 2017) and boom-bust cycles (Bertram & Masel, 2019) can additionally contribute to the maintenance of genetic variation and are probably common in species satisfying these conditions (Kain *et al*., 2015).

Species following such an evolutionary regime should be able to cope also with long term trends as imposed by global change. The next logical step will be to increase the temporal sampling and analyse the spatial dimension. This promises to obtain statistically reliable associations whose functional causalities may then be experimentally validated (but see (Pfenninger & Foucault, 2020)).

## Conclusion

The present study suggested that natural populations can adaptively track the environmental changes in their habitat. It confirms thus findings of a recent experimental study on *Drosophila* (Rudman *et al*., 2021), indicating the relevance of the process for real world populations. Pending further support from additional taxa, these studies provide support for the idea that evolutionary and ecological time scales may not differ at all (Hairston Jr *et al*., 2005). In addition, the rapid and pervasive action of natural selection on the genomic level challenges long-standing population genetic paradigms (Messer *et al*., 2016; Kern & Hahn, 2018).

Beyond these important scientific questions, the presented framework opens the possibility to use population genomic time series for applied functional environmental monitoring (Pfenninger & Bálint, 2022). While currently rather the presence or absence of species is used to evaluate the state of environments, the monitoring of selective changes associated with certain anthropogenic or environmental factors could serve as an early warning system before any community change would be visible. Such knowledge may soon be of crucial importance to direct biodiversity conservation and mitigation strategies in a rapidly changing world.

## Author Contributions

Conceptualization MP, Methodology Development MP, Software Programming MP, Validation MP, Formal Analysis MP, Investigation QF, MP, Resources MP, Data Curation MP, Writing – Original DrMHT Preparation MP, Writing – Review & Editing Preparation QF, MP, Visualization Preparation MP, Supervision MP, Project Administration Management MP, Funding Acquisition MP.

## Acknowledgements

We thank Andreas Wieser for help with sampling and the Abwasserverband Freigericht for generously providing the environmental Hasselbach data. The work was funded by DFG (grant PF390/8-1).

## Data and Code Availability

Sequencing data is publicly available on ENA (project ERP115516, samples ERS4040036-ERS4040041). Core scripts and sync-files were made available on DRYAD (doi.org/10.5061/dryad.fxpnvx0nd).

## SUPPLEMENT

### SUPPL. RESULTS

**Table 1.** Results of two way ANOVA on fitness data of the field population to three experimental temperatures (14°C, 20°C, 26°C) at 5 different sampling dates (Feb. 16, Sept. 16, Feb. 17, Sep. 17, Feb. 18).

| Factor | Sum of sqrs | d.f. | Mean square | F | p (same) |
| --- | --- | --- | --- | --- | --- |
| Sampling date | 0.037 | 4 | 0.009 | 9.509 | 5.07E-06 |
| Exp. Temp | 0.527 | 2 | 0.263 | 269.6 | 1.05E-30 |
| Interaction | 0.028 | 8 | 0.004 | 3.631 | 0.001668 |
| Within | 0.059 | 60 | 0.001 |  |  |
| Total | 0.651 | 74 |  |  |  |

**Table 2.**
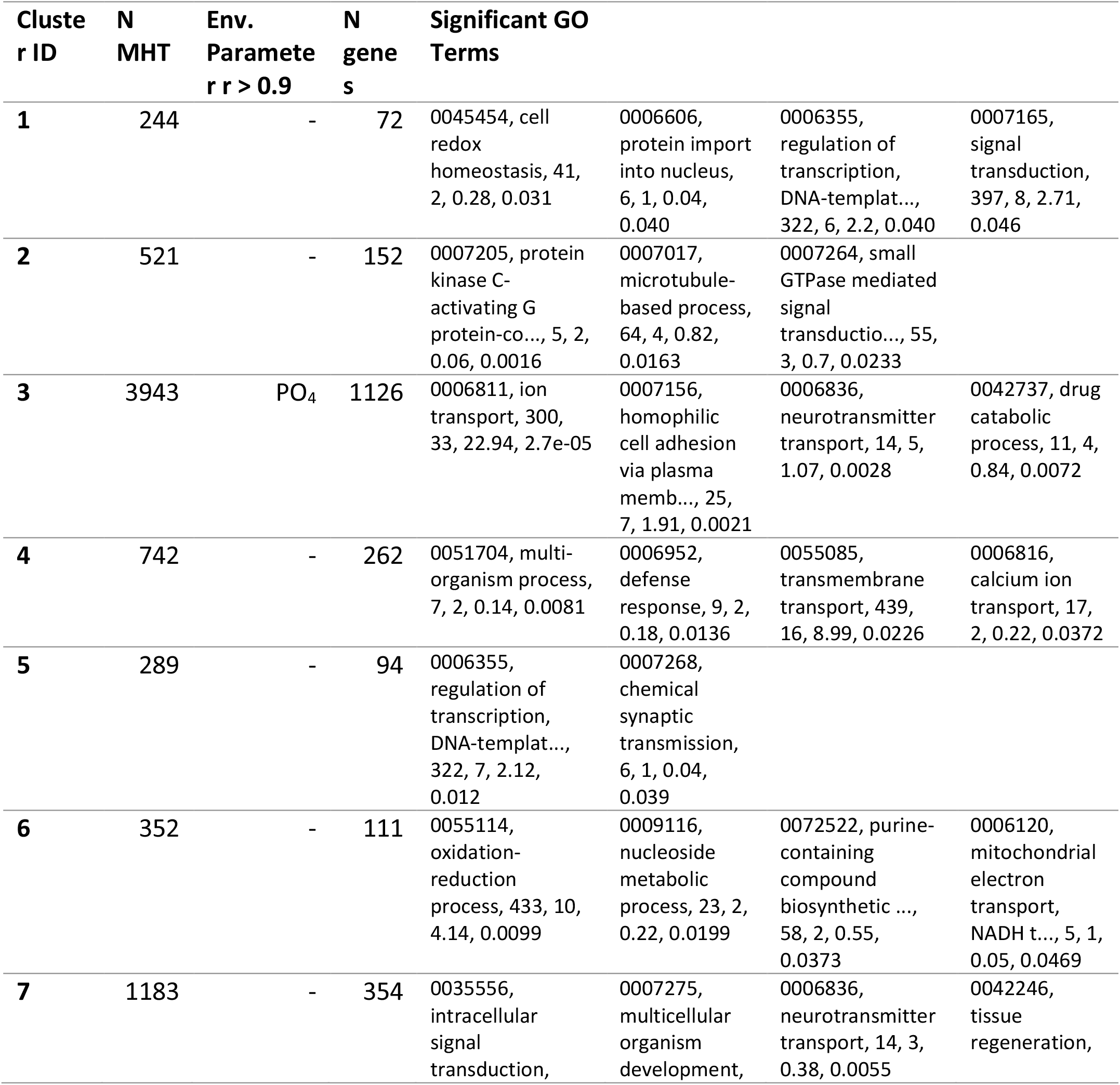
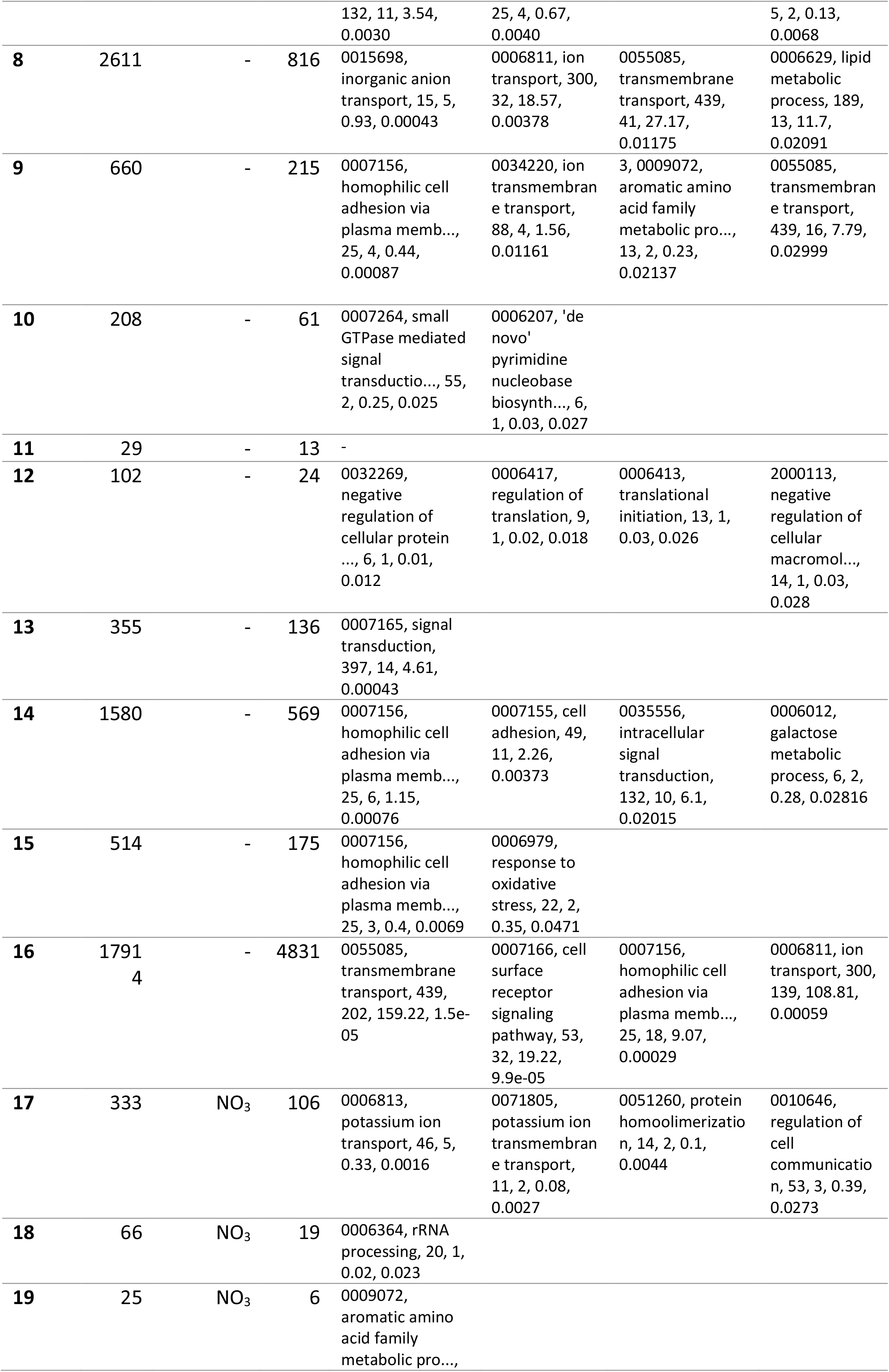
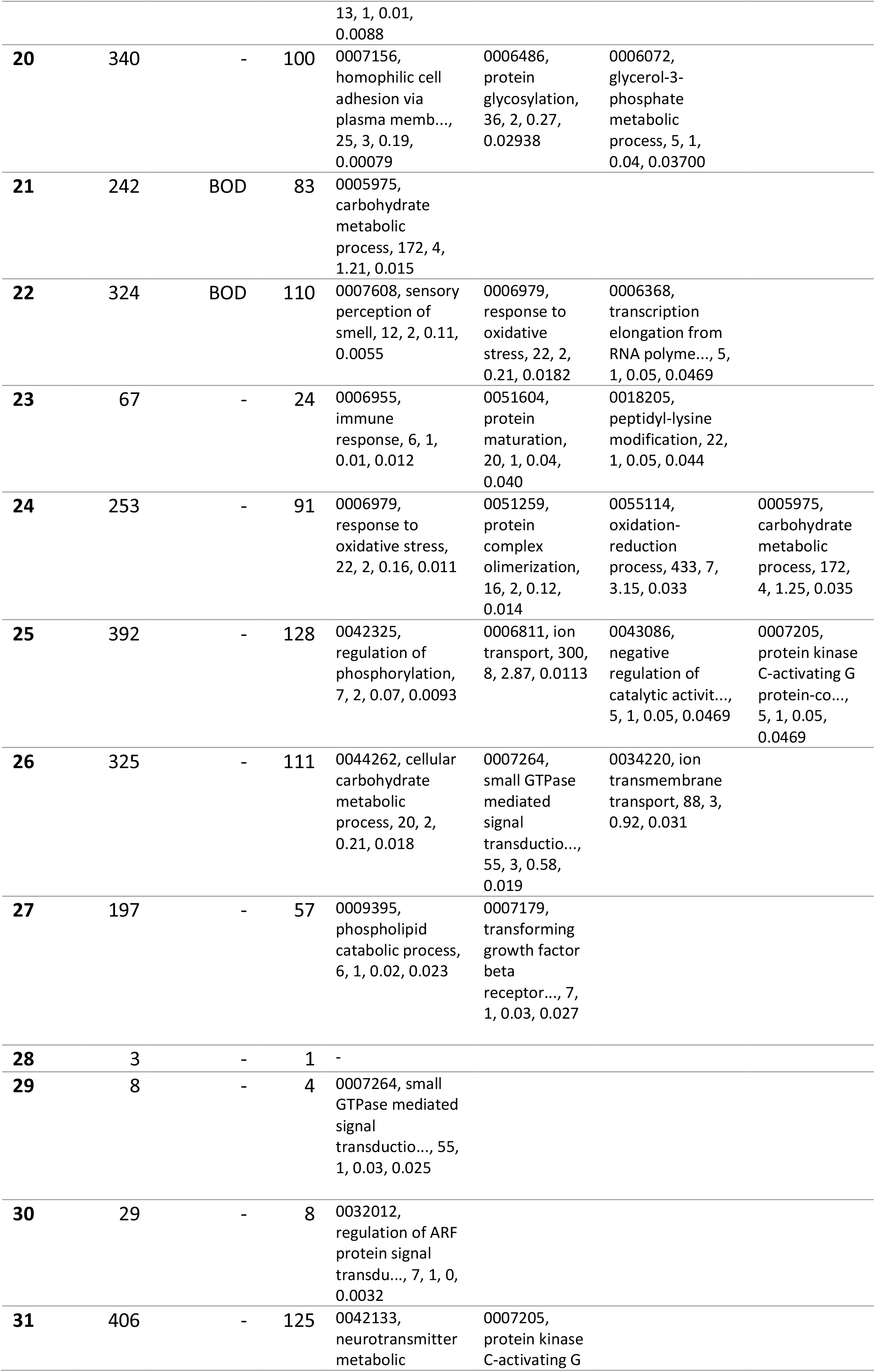
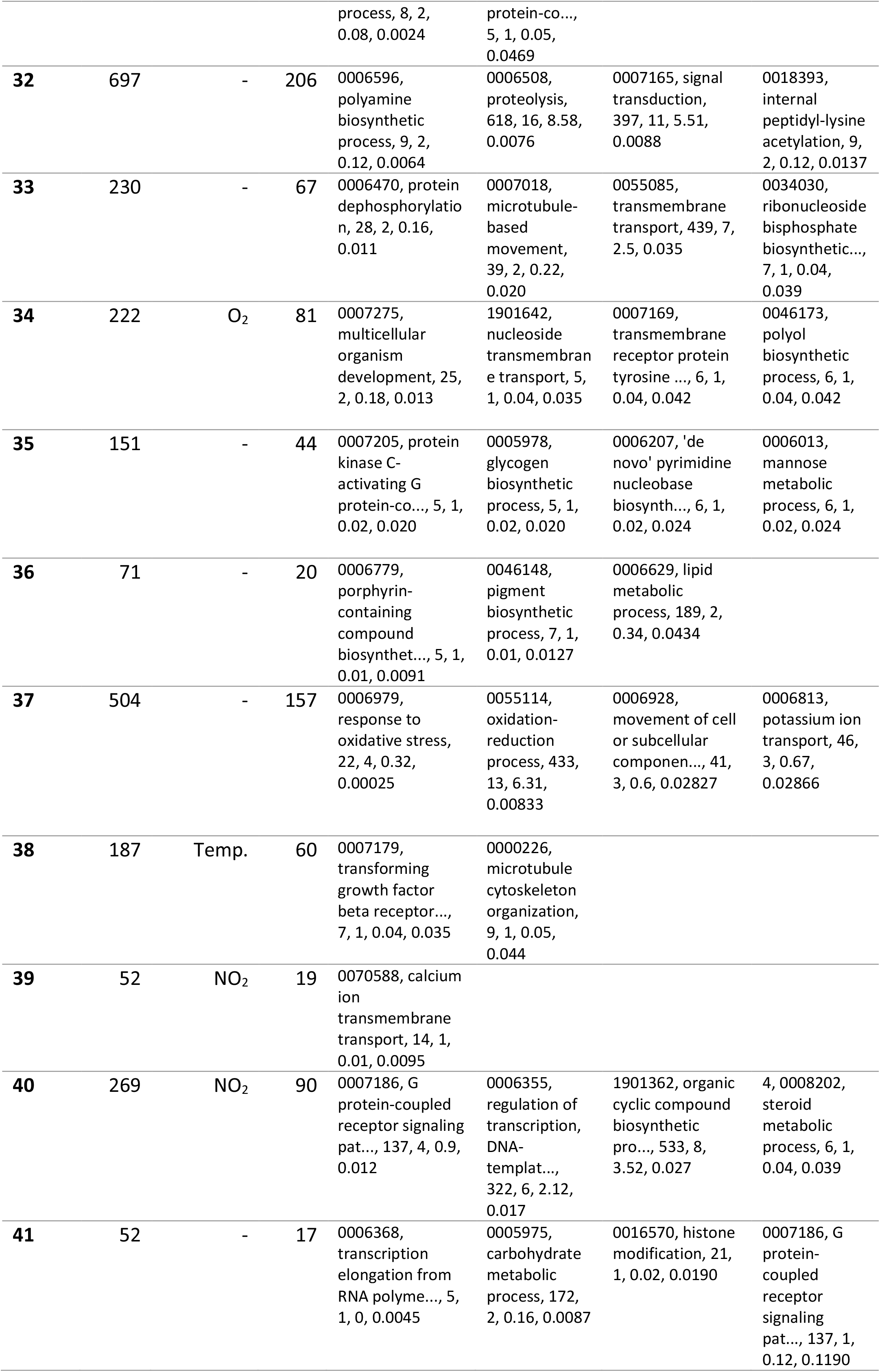
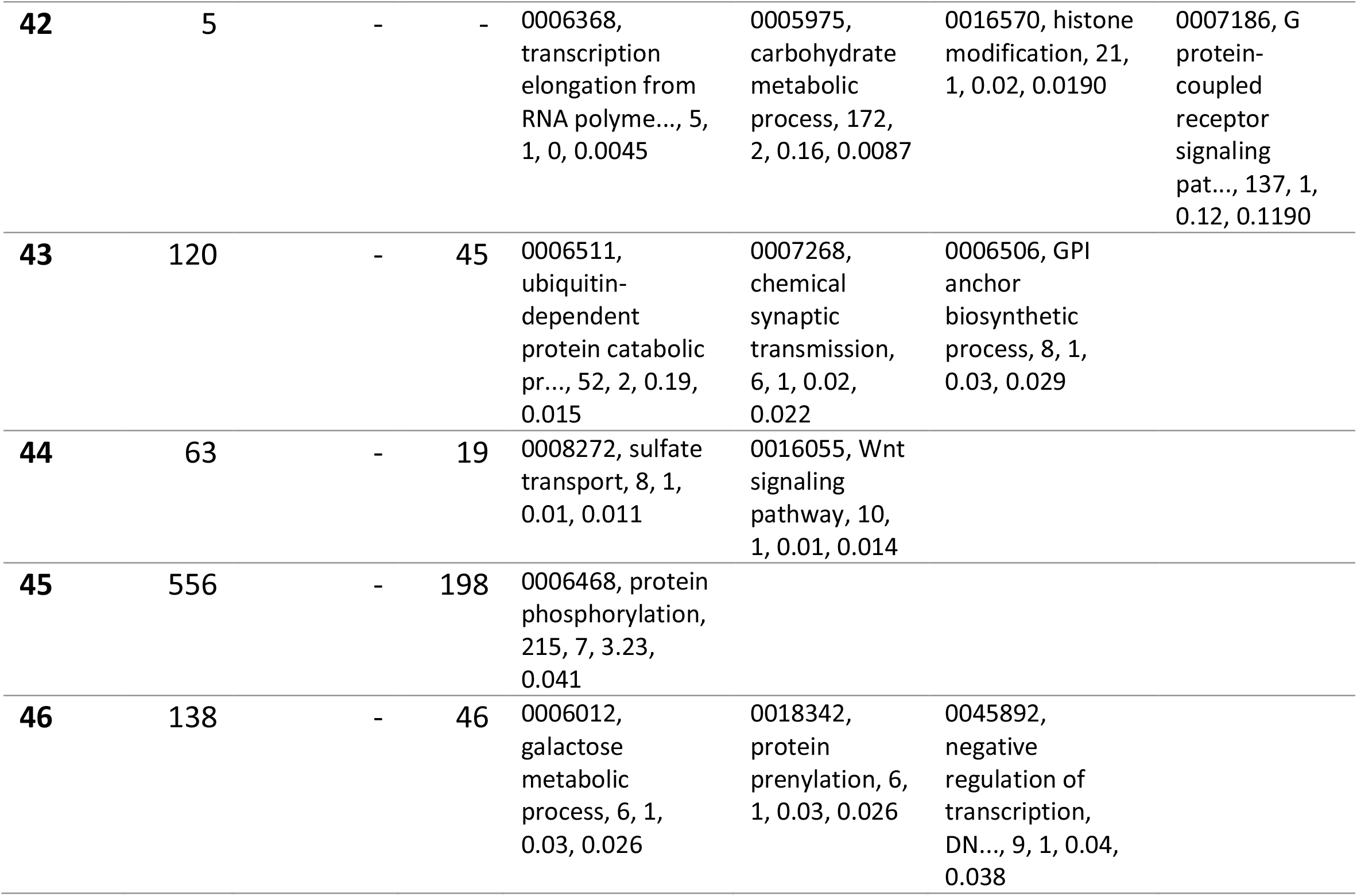
Frequency distribution of MHT cluster patterns, environmental parameter with correlation coefficient > 0.9, number of genes affected and first four most significantly enriched GFO terms for the 46 observed MHT clusters.

| Cluster ID | N MHT | Env. Parameter r r > 0.9 | N genes | Significant GO Terms |
| --- | --- | --- | --- | --- |
| 1 | 244 | - | 72 | 0045454, cell redox homeostasis, 41, 2, 0.28, 0.031<br>0006606, protein import into nucleus, 6, 1, 0.04, 0.040<br>0006355, regulation of transcription, DNA-templat..., 322, 6, 2.2, 0.040<br>0007165, signal transduction, 397, 8, 2.71, 0.046 |
| 2 | 521 | - | 152 | 0007205, protein kinase C-activating G protein-co..., 5, 2, 0.06, 0.0016<br>0007017, microtubule-based process, 64, 4, 0.82, 0.0163<br>0007264, small GTPase mediated signal transductio..., 55, 3, 0.7, 0.0233 |
| 3 | 3943 | PO<sub>4</sub> | 1126 | 0006811, ion transport, 300, 33, 22.94, 2.7e-05<br>0007156, homophilic cell adhesion via plasma memb..., 25, 7, 1.91, 0.0021<br>0006836, neurotransmitter transport, 14, 5, 1.07, 0.0028<br>0042737, drug catabolic process, 11, 4, 0.84, 0.0072 |
| 4 | 742 | - | 262 | 0051704, multi-organism process, 7, 2, 0.14, 0.0081<br>0006952, defense response, 9, 2, 0.18, 0.0136<br>0055085, transmembrane transport, 439, 16, 8.99, 0.0226<br>0006816, calcium ion transport, 17, 2, 0.22, 0.0372 |
| 5 | 289 | - | 94 | 0006355, regulation of transcription, DNA-templat..., 322, 7, 2.12, 0.012<br>0007268, chemical synaptic transmission, 6, 1, 0.04, 0.039 |
| 6 | 352 | - | 111 | 0055114, oxidation-reduction process, 433, 10, 4.14, 0.0099<br>0009116, nucleoside metabolic process, 23, 2, 0.22, 0.0199<br>0072522, purine-containing compound biosynthetic ..., 58, 2, 0.55, 0.0373<br>0006120, mitochondrial electron transport, NADH t..., 5, 1, 0.05, 0.0469 |
| 7 | 1183 | - | 354 | 0035556, intracellular signal transduction,<br>0007275, multicellular organism development,<br>0006836, neurotransmitter transport, 14, 3, 0.38, 0.0055<br>0042246, tissue regeneration, |

|  |  |  |  |  |  |  |  |
| --- | --- | --- | --- | --- | --- | --- | --- |
|  |  |  |  | 132, 11, 3.54,<br>0.0030 | 25, 4, 0.67,<br>0.0040 |  | 5, 2, 0.13,<br>0.0068 |
| **8** | 2611 | - | 816 | 0015698,<br>inorganic anion<br>transport, 15, 5,<br>0.93, 0.00043 | 0006811, ion<br>transport, 300,<br>32, 18.57,<br>0.00378 | 0055085,<br>transmembrane<br>transport, 439,<br>41, 27.17,<br>0.01175 | 0006629, lipid<br>metabolic<br>process, 189,<br>13, 11.7,<br>0.02091 |
| **9** | 660 | - | 215 | 0007156,<br>homophilic cell<br>adhesion via<br>plasma memb...,<br>25, 4, 0.44,<br>0.00087 | 0034220, ion<br>transmembran<br>e transport,<br>88, 4, 1.56,<br>0.01161 | 3, 0009072,<br>aromatic amino<br>acid family<br>metabolic pro...,<br>13, 2, 0.23,<br>0.02137 | 0055085,<br>transmembran<br>e transport,<br>439, 16, 7.79,<br>0.02999 |
| **10** | 208 | - | 61 | 0007264, small<br>GTPase mediated<br>signal<br>transductio..., 55,<br>2, 0.25, 0.025 | 0006207, 'de<br>novo'<br>pyrimidine<br>nucleobase<br>biosynth..., 6,<br>1, 0.03, 0.027 |  |  |
| **11** | 29 | - | 13 | - |  |  |  |
| **12** | 102 | - | 24 | 0032269,<br>negative<br>regulation of<br>cellular protein<br>..., 6, 1, 0.01,<br>0.012 | 0006417,<br>regulation of<br>translation, 9,<br>1, 0.02, 0.018 | 0006413,<br>translational<br>initiation, 13, 1,<br>0.03, 0.026 | 2000113,<br>negative<br>regulation of<br>cellular<br>macromol...,<br>14, 1, 0.03,<br>0.028 |
| **13** | 355 | - | 136 | 0007165, signal<br>transduction,<br>397, 14, 4.61,<br>0.00043 |  |  |  |
| **14** | 1580 | - | 569 | 0007156,<br>homophilic cell<br>adhesion via<br>plasma memb...,<br>25, 6, 1.15,<br>0.00076 | 0007155, cell<br>adhesion, 49,<br>11, 2.26,<br>0.00373 | 0035556,<br>intracellular<br>signal<br>transduction,<br>132, 10, 6.1,<br>0.02015 | 0006012,<br>galactose<br>metabolic<br>process, 6, 2,<br>0.28, 0.02816 |
| **15** | 514 | - | 175 | 0007156,<br>homophilic cell<br>adhesion via<br>plasma memb...,<br>25, 3, 0.4, 0.0069 | 0006979,<br>response to<br>oxidative<br>stress, 22, 2,<br>0.35, 0.0471 |  |  |
| **16** | 1791<br>4 | - | 4831 | 0055085,<br>transmembrane<br>transport, 439,<br>202, 159.22, 1.5e-05 | 0007166, cell<br>surface<br>receptor<br>signaling<br>pathway, 53,<br>32, 19.22,<br>9.9e-05 | 0007156,<br>homophilic cell<br>adhesion via<br>plasma memb...,<br>25, 18, 9.07,<br>0.00029 | 0006811, ion<br>transport, 300,<br>139, 108.81,<br>0.00059 |
| **17** | 333 | NO<sub>3</sub> | 106 | 0006813,<br>potassium ion<br>transport, 46, 5,<br>0.33, 0.0016 | 0071805,<br>potassium ion<br>transmembran<br>e transport,<br>11, 2, 0.08,<br>0.0027 | 0051260, protein<br>homoolimerizatio<br>n, 14, 2, 0.1,<br>0.0044 | 0010646,<br>regulation of<br>cell<br>communicatio<br>n, 53, 3, 0.39,<br>0.0273 |
| **18** | 66 | NO<sub>3</sub> | 19 | 0006364, rRNA<br>processing, 20, 1,<br>0.02, 0.023 |  |  |  |
| **19** | 25 | NO<sub>3</sub> | 6 | 0009072,<br>aromatic amino<br>acid family<br>metabolic pro..., |  |  |  |
|  |  |  |  | 13, 1, 0.01, 0.0088 |  |  |  |
| 20 | 340 | - | 100 | 0007156, homophilic cell adhesion via plasma memb..., 25, 3, 0.19, 0.00079 | 0006486, protein glycosylation, 36, 2, 0.27, 0.02938 | 0006072, glycerol-3-phosphate metabolic process, 5, 1, 0.04, 0.03700 |  |
| 21 | 242 | BOD | 83 | 0005975, carbohydrate metabolic process, 172, 4, 1.21, 0.015 |  |  |  |
| 22 | 324 | BOD | 110 | 0007608, sensory perception of smell, 12, 2, 0.11, 0.0055 | 0006979, response to oxidative stress, 22, 2, 0.21, 0.0182 | 0006368, transcription elongation from RNA polyme..., 5, 1, 0.05, 0.0469 |  |
| 23 | 67 | - | 24 | 0006955, immune response, 6, 1, 0.01, 0.012 | 0051604, protein maturation, 20, 1, 0.04, 0.040 | 0018205, peptidyl-lysine modification, 22, 1, 0.05, 0.044 |  |
| 24 | 253 | - | 91 | 0006979, response to oxidative stress, 22, 2, 0.16, 0.011 | 0051259, protein complex oligomerization, 16, 2, 0.12, 0.014 | 0055114, oxidation-reduction process, 433, 7, 3.15, 0.033 | 0005975, carbohydrate metabolic process, 172, 4, 1.25, 0.035 |
| 25 | 392 | - | 128 | 0042325, regulation of phosphorylation, 7, 2, 0.07, 0.0093 | 0006811, ion transport, 300, 8, 2.87, 0.0113 | 0043086, negative regulation of catalytic activit..., 5, 1, 0.05, 0.0469 | 0007205, protein kinase C-activating G protein-co..., 5, 1, 0.05, 0.0469 |
| 26 | 325 | - | 111 | 0044262, cellular carbohydrate metabolic process, 20, 2, 0.21, 0.018 | 0007264, small GTPase mediated signal transductio..., 55, 3, 0.58, 0.019 | 0034220, ion transmembrane transport, 88, 3, 0.92, 0.031 |  |
| 27 | 197 | - | 57 | 0009395, phospholipid catabolic process, 6, 1, 0.02, 0.023 | 0007179, transforming growth factor beta receptor..., 7, 1, 0.03, 0.027 |  |  |
| 28 | 3 | - | 1 | - |  |  |  |
| 29 | 8 | - | 4 | 0007264, small GTPase mediated signal transductio..., 55, 1, 0.03, 0.025 |  |  |  |
| 30 | 29 | - | 8 | 0032012, regulation of ARF protein signal transdu..., 7, 1, 0, 0.0032 |  |  |  |
| 31 | 406 | - | 125 | 0042133, neurotransmitter metabolic process, 8, 2,<br>0.08, 0.0024 | 0007205, protein kinase C-activating G protein-co...,<br>5, 1, 0.05,<br>0.0469 |  |  |
| **32** | 697 | - | 206 | 0006596,<br>polyamine<br>biosynthetic<br>process, 9, 2,<br>0.12, 0.0064 | 0006508,<br>proteolysis,<br>618, 16, 8.58,<br>0.0076 | 0007165, signal<br>transduction,<br>397, 11, 5.51,<br>0.0088 | 0018393,<br>internal<br>peptidyl-lysine<br>acetylation, 9,<br>2, 0.12, 0.0137 |
| **33** | 230 | - | 67 | 0006470, protein<br>dephosphorylatio<br>n, 28, 2, 0.16,<br>0.011 | 0007018,<br>microtubule-based<br>movement,<br>39, 2, 0.22,<br>0.020 | 0055085,<br>transmembrane<br>transport, 439, 7,<br>2.5, 0.035 | 0034030,<br>ribonucleoside<br>bisphosphate<br>biosynthetic...,<br>7, 1, 0.04,<br>0.039 |
| **34** | 222 | O<sub>2</sub> | 81 | 0007275,<br>multicellular<br>organism<br>development, 25,<br>2, 0.18, 0.013 | 1901642,<br>nucleoside<br>transmembran<br>e transport, 5,<br>1, 0.04, 0.035 | 0007169,<br>transmembrane<br>receptor protein<br>tyrosine ..., 6, 1,<br>0.04, 0.042 | 0046173,<br>polyol<br>biosynthetic<br>process, 6, 1,<br>0.04, 0.042 |
| **35** | 151 | - | 44 | 0007205, protein<br>kinase C-activating G<br>protein-co..., 5, 1,<br>0.02, 0.020 | 0005978,<br>glycogen<br>biosynthetic<br>process, 5, 1,<br>0.02, 0.020 | 0006207, 'de<br>novo' pyrimidine<br>nucleobase<br>biosynth..., 6, 1,<br>0.02, 0.024 | 0006013,<br>mannose<br>metabolic<br>process, 6, 1,<br>0.02, 0.024 |
| **36** | 71 | - | 20 | 0006779,<br>porphyrin-containing<br>compound<br>biosynthes..., 5, 1,<br>0.01, 0.0091 | 0046148,<br>pigment<br>biosynthetic<br>process, 7, 1,<br>0.01, 0.0127 | 0006629, lipid<br>metabolic<br>process, 189, 2,<br>0.34, 0.0434 |  |
| **37** | 504 | - | 157 | 0006979,<br>response to<br>oxidative stress,<br>22, 4, 0.32,<br>0.00025 | 0055114,<br>oxidation-reduction<br>process, 433,<br>13, 6.31,<br>0.00833 | 0006928,<br>movement of cell<br>or subcellular<br>componen..., 41,<br>3, 0.6, 0.02827 | 0006813,<br>potassium ion<br>transport, 46,<br>3, 0.67,<br>0.02866 |
| **38** | 187 | Temp. | 60 | 0007179,<br>transforming<br>growth factor<br>beta receptor...,<br>7, 1, 0.04, 0.035 | 0000226,<br>microtubule<br>cytoskeleton<br>organization,<br>9, 1, 0.05,<br>0.044 |  |  |
| **39** | 52 | NO<sub>2</sub> | 19 | 0070588, calcium<br>ion<br>transmembrane<br>transport, 14, 1,<br>0.01, 0.0095 |  |  |  |
| **40** | 269 | NO<sub>2</sub> | 90 | 0007186, G<br>protein-coupled<br>receptor signaling<br>pat..., 137, 4, 0.9,<br>0.012 | 0006355,<br>regulation of<br>transcription,<br>DNA-templat...,<br>322, 6, 2.12,<br>0.017 | 1901362, organic<br>cyclic compound<br>biosynthetic<br>pro..., 533, 8,<br>3.52, 0.027 | 4, 0008202,<br>steroid<br>metabolic<br>process, 6, 1,<br>0.04, 0.039 |
| **41** | 52 | - | 17 | 0006368,<br>transcription<br>elongation from<br>RNA polyme..., 5,<br>1, 0, 0.0045 | 0005975,<br>carbohydrate<br>metabolic<br>process, 172,<br>2, 0.16, 0.0087 | 0016570, histone<br>modification, 21,<br>1, 0.02, 0.0190 | 0007186, G<br>protein-coupled<br>receptor<br>signaling<br>pat..., 137, 1,<br>0.12, 0.1190 |
| **42** | 5 | - | - | 0006368, transcription elongation from RNA polyme..., 5, 1, 0, 0.0045 | 0005975, carbohydrate metabolic process, 172, 2, 0.16, 0.0087 | 0016570, histone modification, 21, 1, 0.02, 0.0190 | 0007186, G protein-coupled receptor signaling pat..., 137, 1, 0.12, 0.1190 |
| **43** | 120 | - | 45 | 0006511, ubiquitin-dependent protein catabolic pr..., 52, 2, 0.19, 0.015 | 0007268, chemical synaptic transmission, 6, 1, 0.02, 0.022 | 0006506, GPI anchor biosynthetic process, 8, 1, 0.03, 0.029 |  |
| **44** | 63 | - | 19 | 0008272, sulfate transport, 8, 1, 0.01, 0.011 | 0016055, Wnt signaling pathway, 10, 1, 0.01, 0.014 |  |  |
| **45** | 556 | - | 198 | 0006468, protein phosphorylation, 215, 7, 3.23, 0.041 |  |  |  |
| **46** | 138 | - | 46 | 0006012, galactose metabolic process, 6, 1, 0.03, 0.026 | 0018342, protein prenylation, 6, 1, 0.03, 0.026 | 0045892, negative regulation of transcription, DN..., 9, 1, 0.04, 0.038 |  |

**Suppl. Figure 1.**
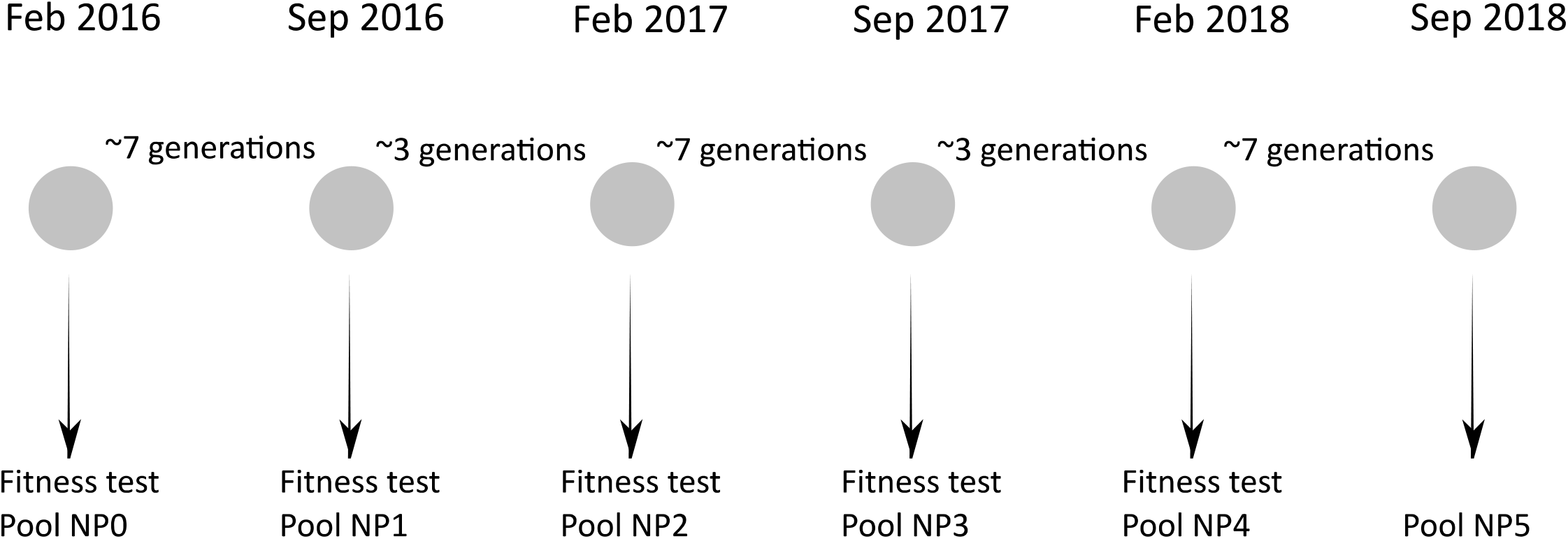
Experimental design. The grey circles indicate the sampling periods.

**Suppl. Fig. 2.**
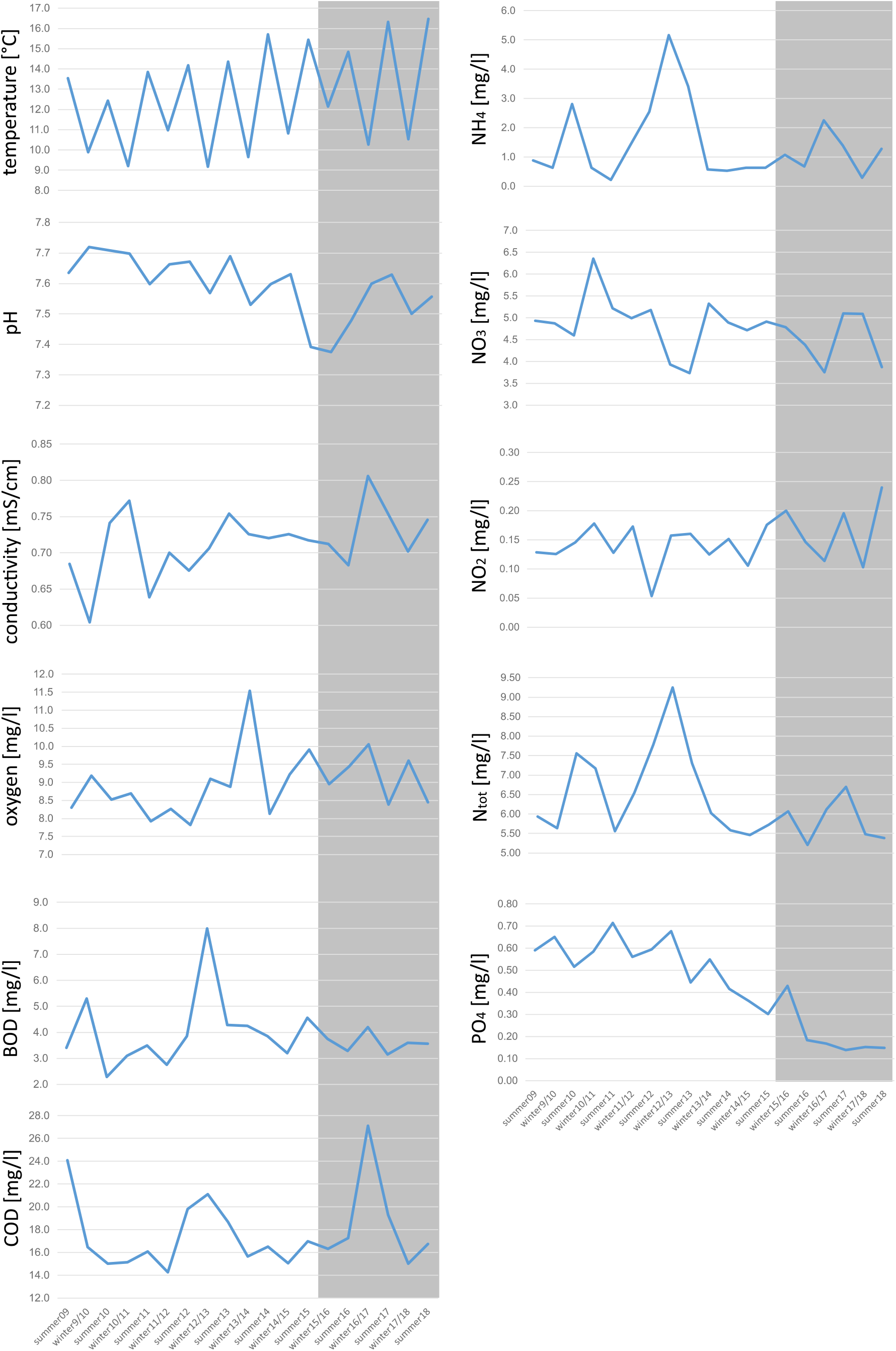
Temporal changes in mean seasonal environmental parameters in the period from 2009-2018 in the Hasselbach. The period relevant for the population genomic analyses is highlighted in grey.

**Suppl. Fig 3.**
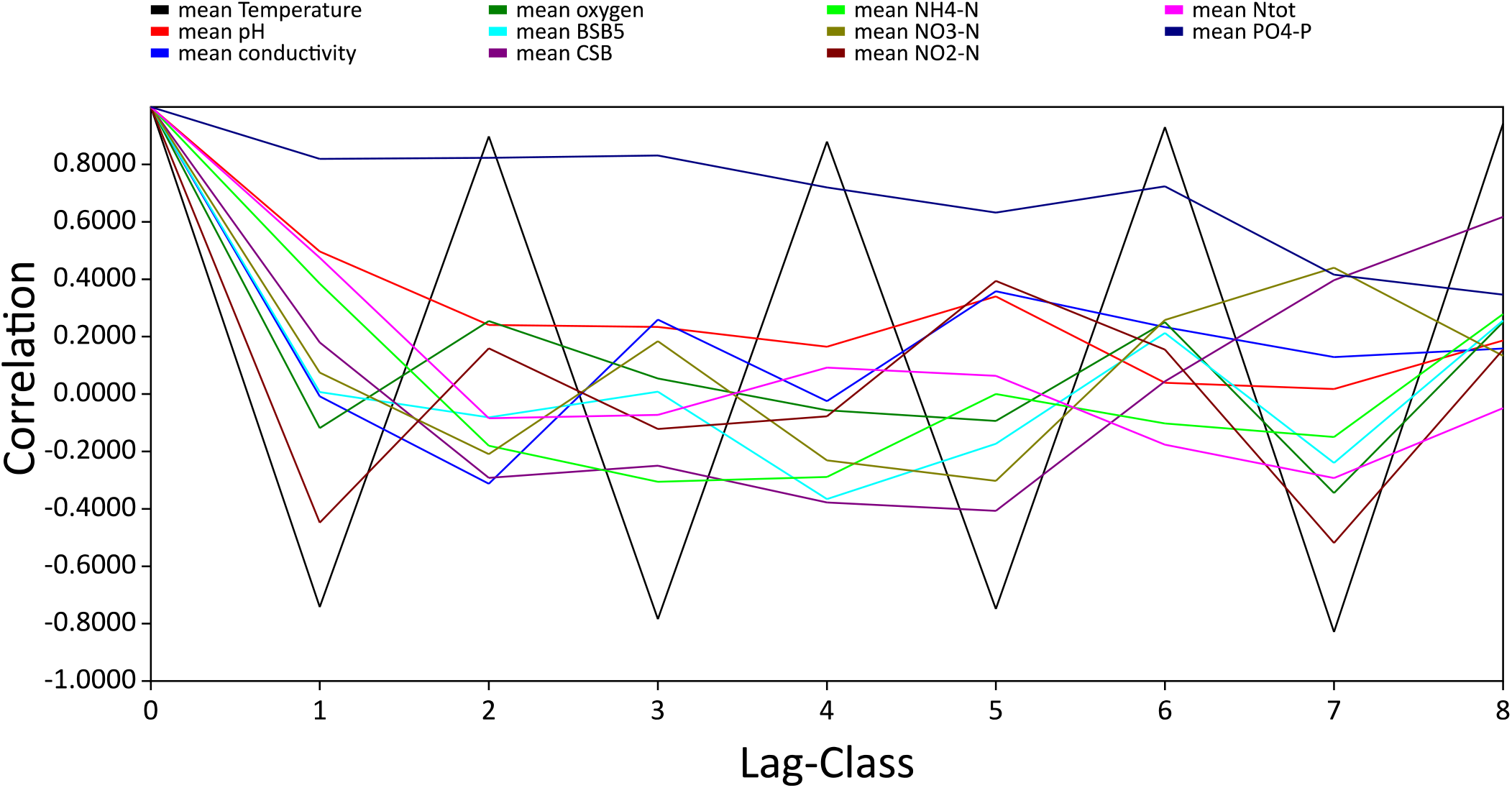
Temporal autocorrelations of environmental parameters.

**Suppl. Fig 4.**
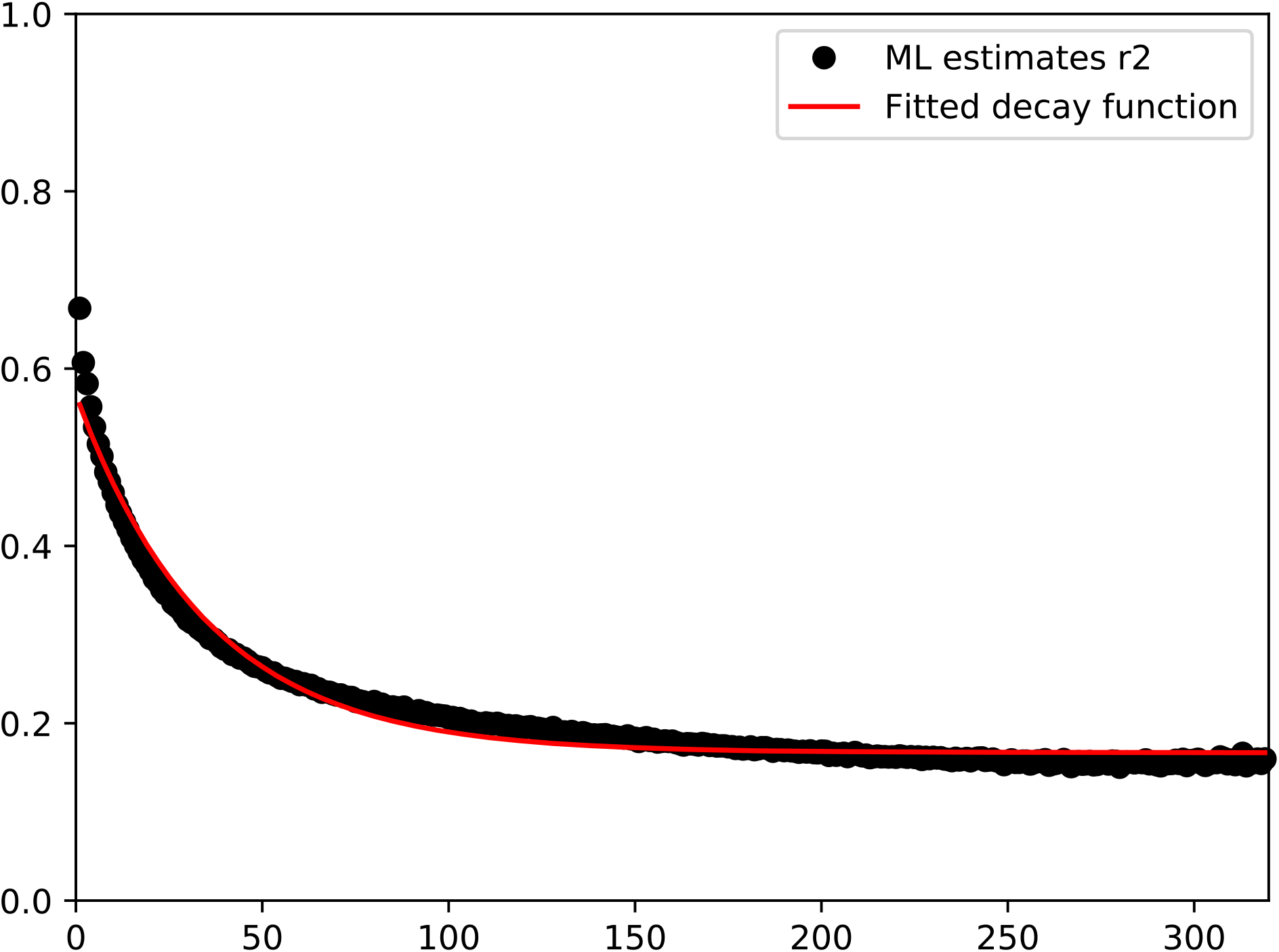
Genome wide mean decay of linkage disequilibrium (r^2^) in dependence of distance in basepairs.

**Suppl. Fig. 5.**
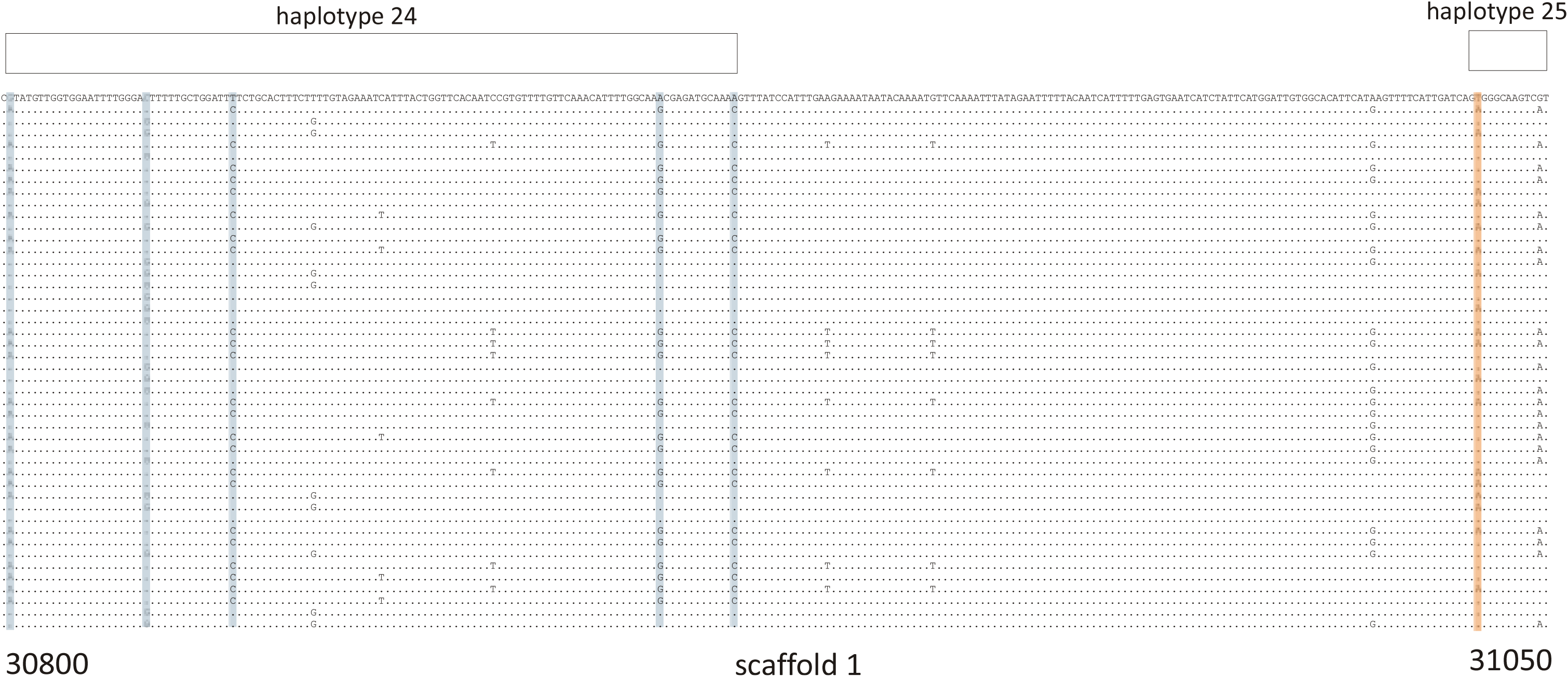
Example for inferred haplotypes of 30 re-sequenced individuals for haplotype 24 and the first SNP of haplotype 25 on scaffold 1 (positions 30800 – 31050). The marker SNPs for haplotype 24 are indicated in blue, for haplotype 25 in orange. The SNPs of haplotype 24 are in full linkage and unlinked to the SNP of haplotype 25.

**Suppl. Fig. 6.**
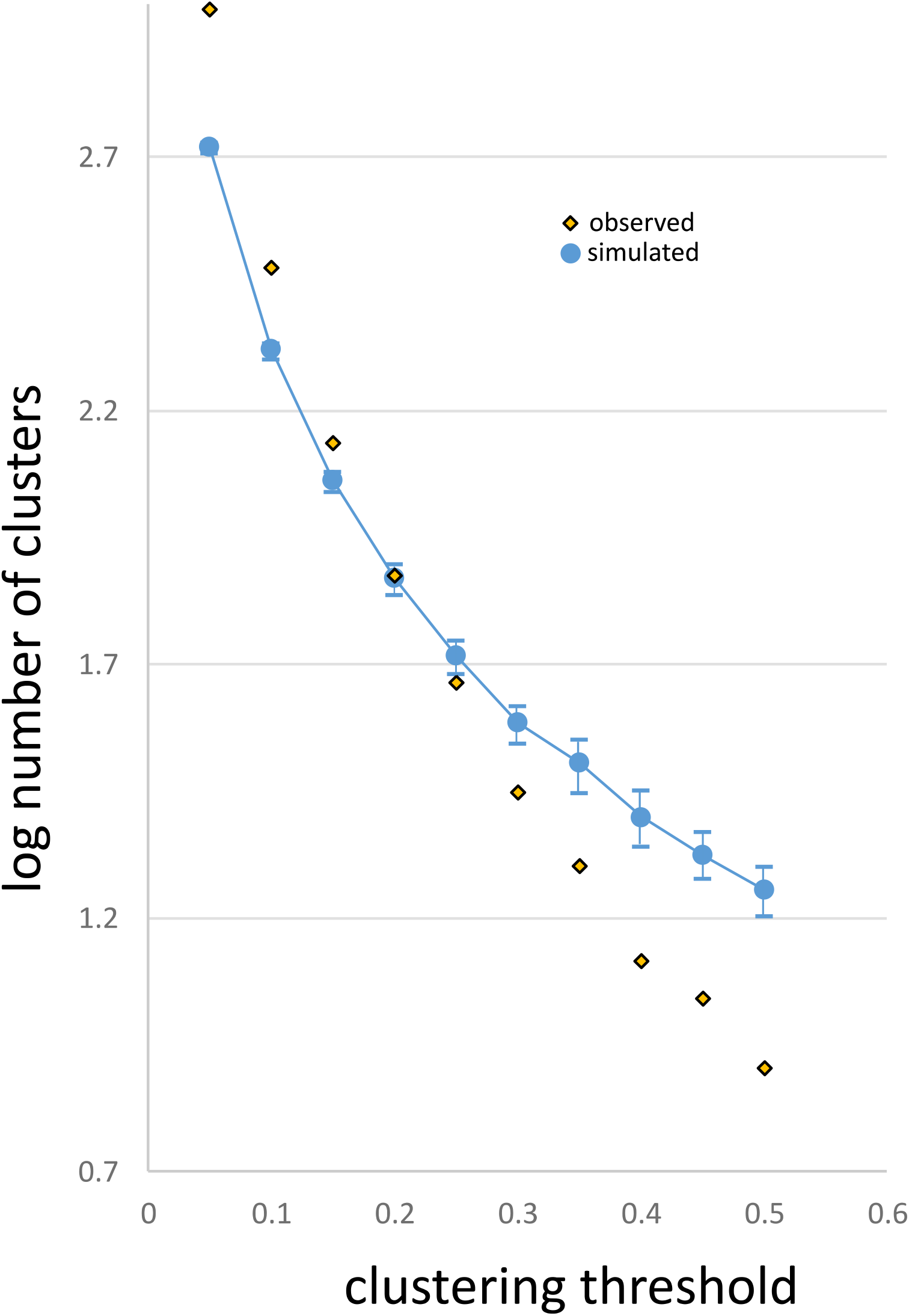
Determination of clustering threshold. Plot of clustering threshold against log number of clusters. In blue, the simulated values with 95% confidence intervals, in yellow with black outline the observed value.

